# Mitochondrial genotype interacts with age and sex, but not nuclear background, to shape locomotory performance across mitonuclear strains of fruit flies

**DOI:** 10.1101/2025.08.22.671777

**Authors:** Rebecca E. Koch, Winston K.W. Yee, Damian K. Dowling

**Affiliations:** Department of Biology, University of Wisconsin-Stevens Point, Stevens Point, WI, 54481, USA; School of Biological Sciences, Monash University, Melbourne, VIC, 3800, Australia

**Keywords:** negative geotaxis, Mother’s Curse, mitonuclear interactions, mtDNA, *Drosophila melanogaster*

## Abstract

The discovery that mitochondrial genomes can harbor functional mutations despite the evolutionarily conserved role of mitochondria has spurred interest in better understanding the ecological and evolutionary consequences of such mitochondrial genetic variation. Because mitochondrial DNA (mtDNA) encodes products that must function in concert with products of the nuclear genome, and because mitochondria are largely maternally inherited, outstanding questions remain as to how, when, and why variation in mtDNA may affect phenotype. In this study, we developed a set of intraspecific “mitonuclear strains” of *Drosophila melanogaster* fruit flies that vary across 13 mtDNA haplotypes and 3 nuclear genetic backgrounds. We created a new apparatus called the Drop Tower to test the performance of these strains in negative geotaxis, a locomotory trait we predicted to be sensitive to variation at the level of mitochondria, across both sexes and two age classes. We found that both mitochondrial and nuclear genetic variation significantly altered how negative geotaxis performance changed with age and across the sexes, though interactions between mitochondrial and nuclear strains did not affect performance. Across most strains, male flies performed more poorly and suffered a steeper decline with age than did females, and young males appeared to vary more in performance across mitochondrial strains. In addition, we discovered a strong effect of small differences of parental age on negative geotaxis— and this effect also varied by mitochondrial strain. Collectively, the results of our study reveal that intraspecific variation in both mitochondrial and nuclear DNA can affect age- and sex-related differences in fruit fly locomotory performance. Further exploring the mechanisms linking these subtle genetic differences to locomotory phenotype and creating differences in response between the sexes will be important to understanding the evolutionary forces shaping the patterns we detected.

## INTRODUCTION

Long thought to be selectively neutral, standing variation in mitochondrial DNA (mtDNA) has been found to influence not only mitochondrial phenotypes, but also key traits important to life-history and fitness, such as lifespan and fertility (Rand *et al*., 2004; Ballard & Pichaud, 2014; Dobler *et al*., 2014; Dowling & Wolff, 2023). That mitochondria have such a core and evolutionarily conserved role at the hub of cellular and physiological processes makes the existence of functional genetic variation that affects mitochondria itself surprising—particularly variation in the mitochondrial genome, which consists of only 13 protein-coding genes in bilaterian metazoans (Blier *et al*., 2001). Because the products of these mtDNA genes must function in concert with nuclear-encoded gene products to form the complexes involved in mitochondrial aerobic respiration, it is important to consider the effects of mitochondrial genetic variation within their nuclear genetic context due to the potential for “mitonuclear” epistatic interactions (Hill *et al*., 2019; Rand & Mossman, 2019). Better understanding the evolutionary ramifications of mitochondrial genetic variation requires teasing apart the contributions of mitochondrial and nuclear gene products (and possible interactions) to mitochondrial and organismal phenotype, as well as the potential evolutionary consequences of mutations in the maternally transmitted mitochondrial genome (Wallace, 2007).

The predominantly maternal transmission of mitochondria complicates the evolutionary context for functional mtDNA variations because without transmission of male mtDNA to their offspring, there is no direct mechanism by which selection can eliminate mutations in mtDNA that harm male function unless these same mutations have similarly negative effects in females; theoretically, this may result in the accumulation of mtDNA mutations that are beneficial or neutral in females, but that have unpredictable or harmful effects to males (Frank & Hurst, 1996; Connallon *et al*., 2018). This possibility, termed the “Mother’s Curse hypothesis” (Gemmell *et al*., 2004), has potential to explain differences in performance between the sexes, such as tendencies for males to live shorter lives than females across many metazoan taxa (Maklakov & Lummaa, 2013; Dowling, 2014). However, how general such Mother’s Curse effects may be across different nuclear genetic contexts and in light of indirect selection on male mtDNA (Unckless & Herren, 2009; Wade & Brandvain, 2009) remains an active area of investigation. Tests of Mother’s Curse effects tend to focus on one of three predictions: that mitochondrial genetic variation will confer greater variation in phenotype in males than in females (the “weak form” of Mother’s Curse); that sexual antagonism will cause a negative intersexual genetic correlation across mitochondrial haplotypes (the “strong form” of Mother’s Curse); or, that experimentally disrupting co-evolved mitochondrial and nuclear genotypes will cause greater fitness loss in males than in females (Dowling & Adrian, 2019). Variation across studies in the approaches used and predictions tested is a likely source of heterogeneity in the outcome of studies testing for evidence of Mother’s Curse effects.

New sets of “mitonuclear” genetic strains that offer variation across both nuclear and mitochondrial genes offer an exciting opportunity to examine the contributions of each—and the epistatic effects between them—to phenotype, as well as whether any sex-biased patterns exist in the expression of mitochondrial genotypic effects. For example, a recent study examined variation in wing size across a nine-by-nine set of mitonuclear genotypes (nine mtDNA haplotypes placed alongside nine different nuclear backgrounds, all sourced from nine allopatric global populations) and discovered evidence not only that mitochondrial and nuclear genes interacted to affect phenotype, but also that levels of phenotypic variance associated with mitochondrial genotypic variation were higher in males than females—supportive of a key prediction of Mother’s Curse (Carnegie *et al*., 2021). However, support for the Mother’s Curse hypothesis remains mixed across studies of different phenotypic traits and genetic lines (e.g. Mossman *et al*., 2016b; a).

In this study, we developed a set of *D. melanogaster* genetic strains that harbored 13 different mitochondrial haplotypes expressed across three different nuclear backgrounds, each created and maintained across two independent duplicate populations. We examined p erformance in a phenotypic trait that is well-studied in *D. melanogaster*: negative geotaxis, or bang-induced climbing. We selected this trait because it comprises multiple aspects of locomotive performance (Grotewiel *et al*., 2005) and has been found to decline with increasing age as well as interact with infection status, oxidative stress, and reductions in superoxide dismutase, a mitochondrial antioxidant (Martin *et al*., 2009; Jordan *et al*., 2012; Linderman *et al*., 2012). Negative geotaxis has recently been used in studies of exercise performance in different *D. melanogaster* mitonuclear strains as well as strains crossing mitochondrial and nuclear types of different *Drosophila* species, finding that both mitochondria and interactions between mitochondrial and nuclear genomes can alter different aspects of response to exercise training (Sujkowski *et al*., 2019; Spierer *et al*., 2021). Further, variation in negative geotaxis performance with age and genetic strain has been found to differ between the sexes (Gargano *et al*., 2005), which suggests that it may be a phenotype sensitive to Mother’s Curse effects as well as functional effects of mitochondrial genetic variation and mitonuclear interactions (Dowling & Adrian, 2019).

Herein, we present the results of 1) a preliminary experiment using a novel negative geotaxis testing apparatus on a set of *D. melanogaster* fruit fly strains that has been previously well studied for functional mitochondrial variation; 2) a main experiment measuring negative geotaxis performance across a new set of mitonuclear genetic strains of fruit flies, across both sexes and two ages; and, 3) a follow-up experiment based on the results of #2, testing for an effect of parental age on negative geotaxis performance across a subset of our strains. Based on previous study of these mitochondrial haplotypes and the negative geotaxis trait, we predicted that both mitochondrial and mitonuclear genetic variation would affect performance across our strains of flies, and that such effects would differ between males and females. Collectively, our study provides insight into how small differences in mitochondrial genotype, nuclear genetic background, age, and even parental age of *D. melanogaster* fruit flies may confer large differences in locomotory phenotype.

## METHODS

This study comprises three separate experiments: a preliminary trial on flies of diverse mitochondrial haplotypes placed against a single nuclear background, a main experiment on a novel, full set of mitonuclear strains created for this study (described below), and a follow-up exploration of the effect of parental age on offspring performance across a subset of these strains (Table 1). All three experiments shared the same fundamental procedures for fly husbandry and negative geotaxis data collection, but differed in the genetic strains used and specifics of analysis.

**Table 1.**
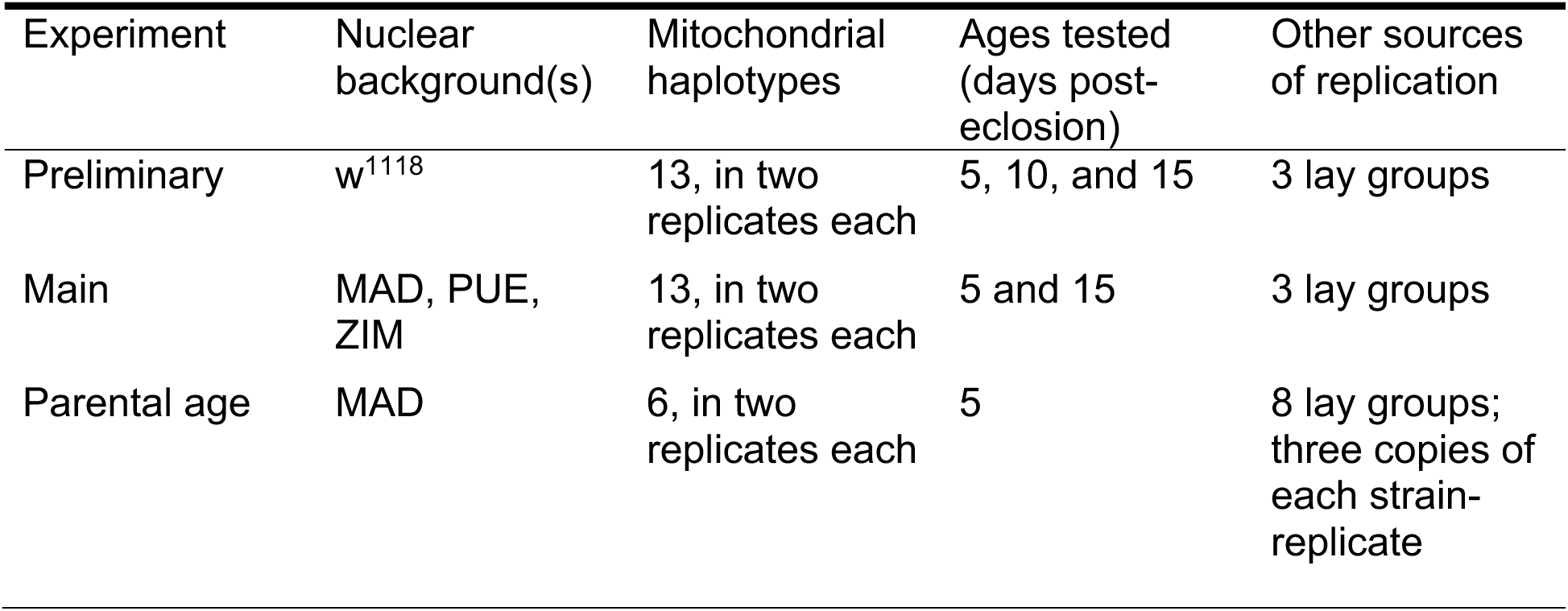
Description of the three experiments that comprise this study.

| Experiment | Nuclear background(s) | Mitochondrial haplotypes | Ages tested (days post-eclosion) | Other sources of replication |
| --- | --- | --- | --- | --- |
| Preliminary | w<sup>1118</sup> | 13, in two replicates each | 5, 10, and 15 | 3 lay groups |
| Main | MAD, PUE, ZIM | 13, in two replicates each | 5 and 15 | 3 lay groups |
| Parental age | MAD | 6, in two replicates each | 5 | 8 lay groups; three copies of each strain-replicate |

All flies throughout this experiment were maintained on a medium of yeast, dextrose, agar, and potato (with 12 mL/L nipagen and 5 mL/L propionic acid) contained in 40 mL plastic vials and sprinkled with granules of live yeast to provide *ad lib* access. Flies were kept on a 12 hour light-dark cycle under constant 25 °C temperature control.

### Negative geotaxis testing apparatus and data collection

As the procedure for inducing and recording negative geotaxis response is identical for all three experiments, we describe it prior to discussing the details that distinguish each experiment. Many different methods exist for testing negative geotaxis response in fruit flies; we chose to emulate a procedure akin to “rapid iterative negative geotaxis” as described in Gargano *et al*. (2005). We designed a new apparatus intended to standardize the timing and magnitude of each induction of negative geotaxis by dropping vials of flies against a firm surface. To construct the “Drop Tower” apparatus, we fastened six 60 cm polyvinyl chloride pipes (3 cm diameter) together and glued them to a stiff posterboard that we mounted above a laboratory bench (Figure 1). 50 cm above the bench surface, we cut a slit in the outward-facing portion of each pipe such that we could slide a painter’s spackling blade (25 cm wide) into the slit, effectively closing all six pipes. We could then load vials of experimental flies into the top of each pipe above the blade, then rapidly pull the blade out from the pipes to release the vials of flies to fall 50 cm to the bench top below (Supplementary Video). We placed one layer of removable mounting adhesive (Bostik Blu Tack) at the point of contact on the bench surface, under the Drop Tower, such that vials would stick rather than bounce upon impact. We mounted the pipes such that their ends were 6.5 cm above the bench surface so that almost the entirety of all dropped vials would be visible, but the top most portions would still be inside the pipes to keep the vials from toppling (Supplementary Video). Flies in vials released through the Drop Tower are knocked to the bottoms of their vials upon impact with the benchtop, inducing a negative geotaxis climbing response. Six vials of flies could be tested (“dropped”) simultaneously across the six pipes of the Drop Tower. We tested groups of six vials in a series of consecutive drops; vials of flies were loaded into the Drop Tower and dropped, allowed to climb for 6-10 seconds, then immediately loaded and dropped again for up to five total consecutive repetitions. We rotated the vials across the different pipes of the Drop Tower between repetitions to control for any variability among pipes, and we altered which genetic strains were included within each group of six vials dropped together across different trials. We transferred each vial of flies from their media-filled housing vials into empty vials sealed with plastic caps immediately before dropping.

**Figure 1.**
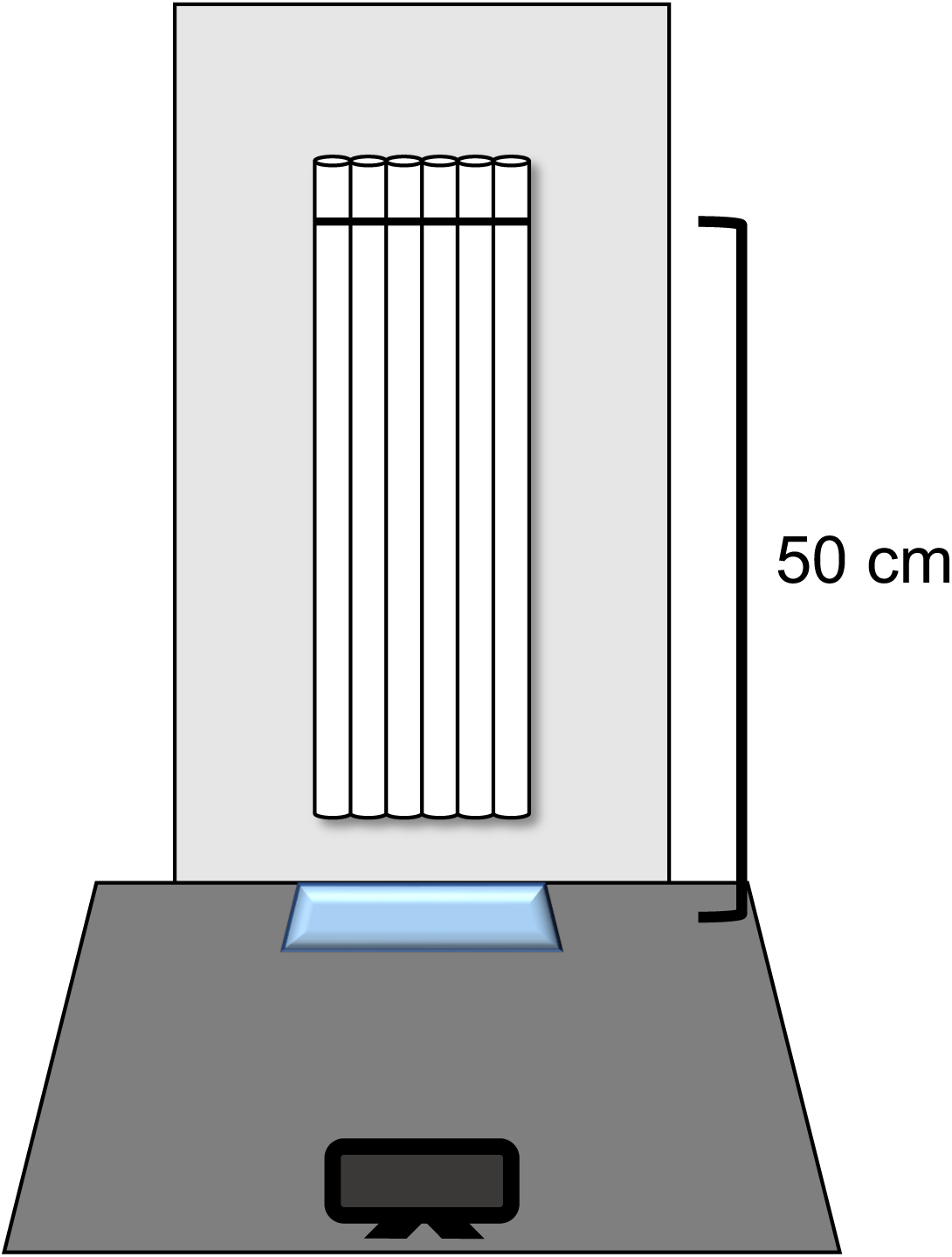
The Drop Tower, a new apparatus built for testing negative geotaxis performance. Six vials containing fruit flies are dropped simultaneously though plastic pipes (white) on to a sticky surface of mounting adhesive (blue), and the resulting climbing response is captured on video (black).

During experimental trials, we used an iPhone 6 (Apple, Inc.) mounted 25 cm from the apparatus (Figure 1) to record the impact of the vials and the subsequent climbing of flies, using a slow-motion setting to record at 240 frames per second. We also placed a timer showing time to the hundredth of a second within the camera’s view. From the video recordings of each trial, we referenced this timer to pause the video exactly four seconds after the vials impacted, at which point many of the flies in each vial would be climbing. From this still image, we extracted data on performance in each vial (see below for specifics for each experiment).

### Preliminary trials on white-eyed fruit flies

The Dowling Lab has maintained populations of white-eyed *Drosophila melanogaster* that are near-isogenic for the *w^1118^* nuclear genotype (BloomingtonStock Number 5905), but that differ in their mitochondrial haplotype. These haplotypes were originally sourced from *D. melanogaster* populations worldwide, representing a large breadth of the mitochondrial genetic variation present in this species (Wolff *et al*., 2016). The Dowling Lab acquired copies of these strains in 2008 from David Clancy (Lancaster University), at which point each strain was split into two parallel populations of the same mitochondrial haplotype; we will refer these duplicate populations as “strain replicates,” as they serve to provide biological replication of each mitochondrial haplotype (Camus *et al*., 2012; Nagarajan-Radha *et al*., 2020). In the years since, the strains have been maintained as near-isogenic for the nuclear genome through continual backcrossing of strain females with inbred *w^1118^* males (themselves maintained as near-isogenic through full-sibling propagation; Innocenti *et al*., 2011; Camus *et al*., 2012). The haplotypes represented in these strains were originally sourced from: Alstonville, New South Wales, Australia (ALS); Barcelona, Spain (BAR); Brownsville, Texas, USA (BRO); Dahomey, Benin (DAH); Hawaii, USA (HAW); Israel (ISR); Japan (JAP); Madang, Papua New Guinea (MAD); Mysore, India (MYS); Oregon, USA (ORE); Puerto Montt, Chile (PUE); Sweden (SWE); and Zimbabwe (ZIM). We first used these 13 strains (in 26 total strain replicates, all in the *w^1118^* nuclear background) to perform a preliminary study to pilot our methods.

In June 2018, we placed five pairs of adult flies (5 days post-eclosion in age) of each strain replicate in vials containing media to mate and enable females lay eggs for 24 hours; pairs were transferred to fresh vials of medium each day for four days to provide four sets (“lay groups”) of potential offspring to collect for experimentation.

Three lay groups—those from eggs laid on the first, third, and fourth day—were used in this experiment. We collected up to 15 virgin male and female flies of each strain replicate from each lay group for experimental testing. After collection as virgins, experimental flies entered a schedule in which they were transferred to fresh vials of medium every 2 or 3 days, and tested in the Drop Tower every 5 days. The three lay groups were collected, transferred, and tested from 15 June to 13 July 2018 such that each vial of flies was tested at ages 5, 10, 15, 20, and 25 days since eclosion, though only ages 5, 10, and 15 were analysed because performance decreased dramatically by day 15 (see below) and few flies climbed at all by the age of 20 days.

In each day of running a negative geotaxis trial, we tested one vial of 5 to 15 flies per experimental unit (i.e. one vial of each sex per mitochondrial strain replicate; Table 1). With 26 strain replicates and two sexes, this is a total of 52 vials, dropped across 9 groups of 4-6 vials; these trials took about 45 minutes in total, and began around 8:30 AM (90 minutes after the start of the flies’ daily light cycle). After testing, flies were returned to fresh vials of medium and maintained as described above until the next testing period or the end of the experiment.

Extracting data from each trial was a two-step process: first, we used the video recordings to count any flies that had climbed out of view at the top of each vial, and the flies that had never climbed (i.e. were still resting at the vial bottom); second, we collected and assessed a screen-capture of the video paused exactly four seconds after vial impact. From this screen-captured image, we cropped out the central regions of the six vials (a region sized 19 cm wide by 6 cm tall) such that only climbing flies were visible. We then ran these cropped images through a macro in ImageJ (Rasband, 1997) that used particle detection to measure the height climbed (position along the y-axis) by each fly (Gargano *et al*., 2005), which we then matched to a specific vial using position along the x-axis. Finally, we used these two sources of data to calculate the negative geotaxis performance of each drop of each vial as the average height climbed by the flies in that vial; flies that never climbed were included in the calculation as values of zero, and flies that climbed to the top of the vial within the four-second time frame were given the maximum vial height of 6.5 cm. Each vial was dropped five consecutive times per trial, though we analyzed only the first three drops in this preliminary experiment.

To assess the results of this preliminary study, we fit a linear mixed effects model using the lme4 package (v. 1.1-37; Bates, 2014) in R (v. 4.5.1; R Core Team, 2025). This model included: average negative geotaxis response of each vial (i.e. average cm climbed) as the response variable; fixed effects of age (3 levels), sex (2 levels), mitochondrial strain (13 levels), and all possible two-way interactions; random intercept effects to account for any overall variation across lay group (3 levels), order dropped within a trial (“drop order”; 9 levels), a composite variable of lay group, age, and drop order (“drop group,” to account for the specific group of six vials dropped together; 3 lay groups x 3 ages x 9 drop order levels = 81 levels), the drop number (representing the three repeated measures; 3 levels), strain replicate (13 strains x 2 replicates = 26 levels), and vial (26 strain replicates x 2 sexes x 3 ages = 156 levels); and, a random slope term allowing the effects of age, sex, and their interaction to vary across strain replicates (i.e. “Age + Sex + Age:Sex | Strain Replicate”), and a random slope term allowing the effects of drop number and age to vary across vials (accounting for the three repeated measures). We set the sum of contrasts to zero, and evaluated the significance of fixed effects using the anova function in the lmerTest package (v. 3.103; Kuznetsova *et al*., 2017), which performs F tests on fixed effects using Satterthwaite approximation of degrees of freedom. We used the drop1 function with Kenward-Rogers approximation to evaluate potential fixed effects to drop from the full model, resulting in dropping two-way interactions involving sex from the model. Finally, we reduced the complexity of our random effects statements in a stepwise fashion until the model converged on estimates without singularity warnings. We first simplified the random effects by converting random slope terms to random intercept terms containing an interaction of the focal variables (e.g. “Age | Strain Replicate” becomes “1 | Age : Strain Replicate”), which reduces complexity while retaining some of the full model’s structure. We then further simplified or dropped random effects terms that explained little to no variance until the model reached convergence. We did not simplify models past the point of convergence in order to avoid inflating denominator degrees of freedom in our fixed effects due to dropping terms important to the replication structure of our dataset (Arnqvist, 2020). Our final model includes fixed effects of age, sex, mitochondrial strain, and a two-way interaction of age and mitochondrial strain; and, random intercept effects of drop group, lay group, drop number, the interaction of vial and drop number, and the interactions of mitochondrial strain replicate with age, with sex, or with both age and sex (Table 2).

**Table 2.**
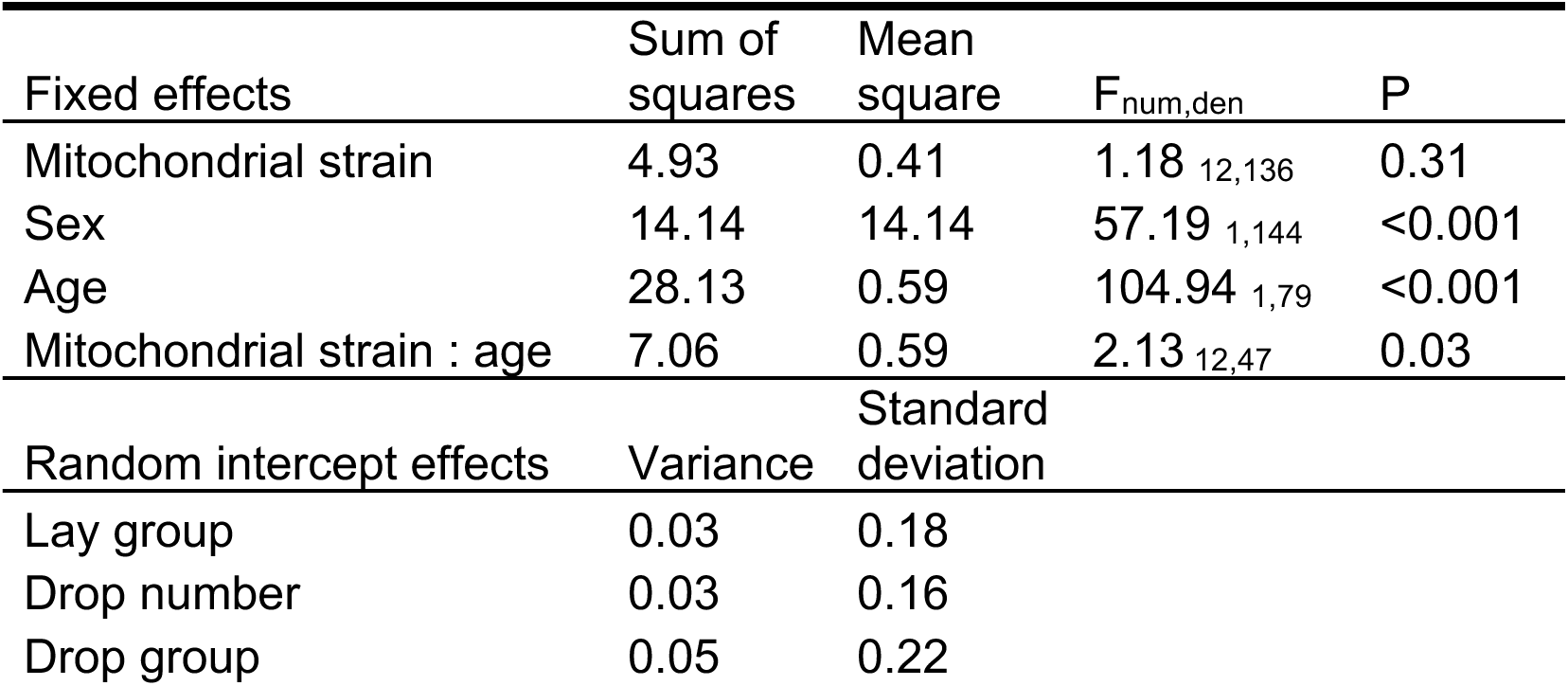
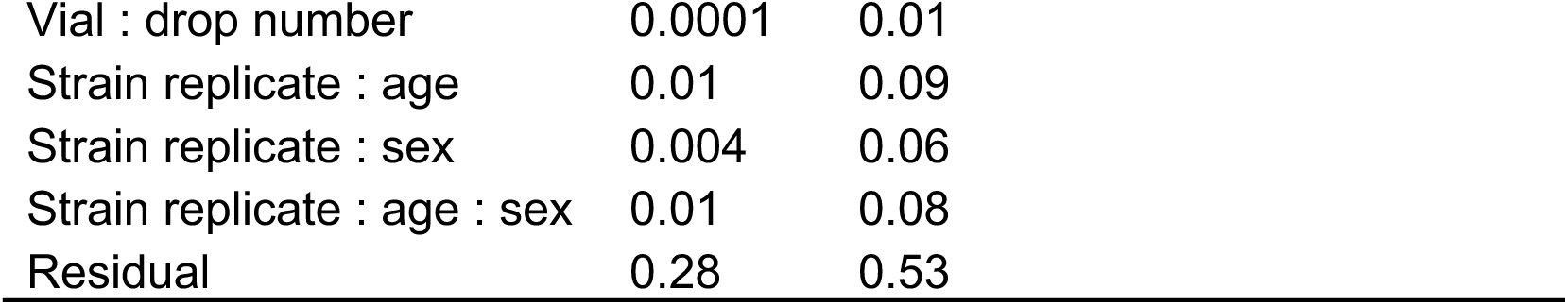
Results of the final, reduced linear mixed effects model testing for the effects of focal fly age, sex, and mitochondrial strain on negative geotaxis performance, across 13 replicated mitochondrial strains within the *w^1118^* nuclear background. In this experiment, we tested the same groups of flies across three ages: 5, 10, and 15 days post-eclosion. Subscript values on the F statistic refer to the numerator and denominator degrees of freedom, respectively, approximated with Satterthwaite’s method.

| Fixed effects | Sum of squares | Mean square | F<sub>num,den</sub> | P |
| --- | --- | --- | --- | --- |
| Mitochondrial strain | 4.93 | 0.41 | 1.18<sub>12,136</sub> | 0.31 |
| Sex | 14.14 | 14.14 | 57.19<sub>1,144</sub> | <0.001 |
| Age | 28.13 | 0.59 | 104.94<sub>1,79</sub> | <0.001 |
| Mitochondrial strain : age | 7.06 | 0.59 | 2.13<sub>12,47</sub> | 0.03 |
| Random intercept effects | Variance | Standard deviation |  |  |
| Lay group | 0.03 | 0.18 |  |  |
| Drop number | 0.03 | 0.16 |  |  |
| Drop group | 0.05 | 0.22 |  |  |

|  |  |  |
| --- | --- | --- |
| Vial : drop number | 0.0001 | 0.01 |
| Strain replicate : age | 0.01 | 0.09 |
| Strain replicate : sex | 0.004 | 0.06 |
| Strain replicate : age : sex | 0.01 | 0.08 |
| Residual | 0.28 | 0.53 |

### Main experiment on mitonuclear strains

Starting in January 2018, we used backcrossing to introgress the mitochondrial haplotype of each of the 13 strains described above into three new nuclear backgrounds, originally sourced from Madang, Papua New Guinea (MAD), Puerto Montt, Chile (PUE), and Zimbabwe (ZIM)— lines from which three of the mitochondrial haplotypes of our strains were derived. In the first generation, we sourced five males from each chosen nuclear background (maintained as near-isogenic populations through over 20 generations full-sibling crossing) and paired them with five virgin females from the *w^1118^* strain replicates; each subsequent generation, we crossed males of each target nuclear background with virgin females of each strain produced from the previous generation of backcrossing. The end result is a new set of 39 mitonuclear strains (3 nuclear backgrounds x 13 mitochondrial haplotypes), created and maintained in duplicated populations for a total of 78 strain replicates. At the time of the experiment, the PUE strains had undergone 12 generations of introgressive backcrossing, and the MAD and ZIM strains had undergone 15 generations, which in theory would have replaced at least around 99.97% of the original *w^1118^* nuclear background with nuclear alleles from each of the MAD, PUE and ZIM backgrounds.

In October 2018, we first collected the parents of experimental flies as five pairs of virgin flies per strain replicate. Starting with one-day-post-eclosion virgins, we transferred flies into new vials of fresh media every 24 hours for three days, retaining the three resultant sets of vials to form three lay groups of the focal generation of flies that differed only in the age of their parents at time of laying (ages 1-2, 2-3, or 3-4 days). Each time that we transferred the parental flies out of a vial, we carefully used a metal spatula to trim off excess eggs from vials that contained more than around 80 eggs. Upon the eclosion of flies of each lay group, we collected one vial of 15 virgin female flies and one vial of 15 virgin male flies from each strain replicate for testing. In total, each of the 78 strain replicates was represented by six vials of 15 flies: one vial per sex, per lay group. The only exception is for the BRO strain in the MAD nuclear background, which proved to be almost entirely infertile for crosses within the strain (vs. with backcrossing, as during maintenance), so we were unable to collect sufficient flies for testing in this instance; subsequent study revealed that flies of this particular mitonuclear strain have particularly poor pre-copulatory mating success (Koch & Dowling, 2022), providing a possible explanation for this observation (also see Clancy *et al*., 2011).

Once collected, we transferred experimental flies to vials containing fresh media on days 2, 5, 7, 10, and 12 post-eclosion, and tested flies in negative geotaxis trials on days 5 and 15. Based on results from the preliminary study on *w^1118^* flies, we anticipated that these two time points would capture variation in age-related decline in performance. We monitored numbers of flies that died prior to experimentation such that we had an accurate count of the numbers of flies tested in each vial in every trial. Data extraction from the videos of these trials was therefore nearly identical to that in the preliminary experiment, except that we collected data on all five repeated measures per vial, rather than only the first three. Otherwise, we again counted flies that reached the very tops of their vials from video, we used our macro in ImageJ to measure the heights climbed by flies mid-way up the vial sides after four seconds, and we could calculate the numbers of flies that did not climb within the trial. From this data, we calculated average height climbed by the flies contained within each vial for each negative geotaxis drop event, resulting in five repeated measures per vial.

Our statistical analysis of this data resembles that of our preliminary study, but also accounts for variation in nuclear background. Using the lme4 package, we first ran a full model that featured: a response variable of average height climbed by flies in a vial; fixed effects of nuclear background (3 levels), mitochondrial strain (13 levels), sex (2 levels), age (2 levels), and all possible two- and three-way interactions between them; random intercept effects to account for variation across lay group (3 levels), order dropped within a trial (26 levels), a composite variable of lay group, age, and drop order (“drop group,” accounting for the specific group of six vials dropped together; 156 levels), and the drop number (representing the five repeated measures; 5 levels); a random slope term allowing the effects of age, sex, and their interaction to vary across strain replicate; and, a random slope term allowing the effects of drop number and age to vary across vials to account for the repeated measures of each vial. Again, we set sum of contrasts to zero, evaluated the significance of fixed effects using the anova function in the lmerTest package, and used the drop1 function as described above to inform model reduction, which resulted in dropping all three-way interactions and, notably, the two-way interaction between mitochondrial strain and nuclear background. Our final model included fixed effects of: age, sex, nuclear background, and mitochondrial strain; five two-way interactions (between nuclear background and age or sex, between mitochondrial strain and age or sex, and between age and sex); and, the three-way interaction between mitochondrial strain, sex, and age. This final model also included random intercept effects of drop group, drop number, drop order, lay group, the interaction of vial and drop number, the two-way interaction of strain replicate and age, and the three-way interaction of strain replicate with age and sex (Table 3).

**Table 3.** Results of the final model testing for the effects of focal fly age, sex, nuclear background, and mitochondrial strain on negative geotaxis performance in fruit flies across our main set of strains. These strains comprised 13 replicated mitochondrial strains across three nuclear backgrounds, each measured at two ages (5 and 15 days post-eclosion). Subscript values on the F statistic refer to the numerator and denominator degrees of freedom, respectively, approximated with Satterthwaite’s method.

| Fixed effects | Sum of squares | Mean square | F<sub>num, den</sub> | P |
| --- | --- | --- | --- | --- |
| Nuclear background | 0.48 | 0.24 | 0.63<sub>2,204</sub> | 0.54 |
| Mitochondrial strain | 1.57 | 0.13 | 0.34<sub>12,113</sub> | 0.98 |
| Sex | <0.001 | <0.001 | 0.0003<sub>1,119</sub> | 0.99 |
| Age | 5.43 | 5.43 | 14.23<sub>1,150</sub> | <0.001 |
| Nuclear background : sex | 11.05 | 5.53 | 14.49<sub>2,131</sub> | <0.001 |
| Nuclear background : age | 4.35 | 2.18 | 5.71<sub>2,203</sub> | 0.004 |
| Mitochondrial strain : sex | 8.04 | 0.67 | 1.76<sub>12,119</sub> | 0.06 |
| Mitochondrial strain: age | 1.78 | 0.15 | 0.39<sub>12,111</sub> | 0.96 |
| Sex : age | 41.65 | 41.65 | 109<sub>1,117</sub> | <0.001 |
| Mitochondrial strain : sex : age | 9.50 | 0.79 | 2.08<sub>12,117</sub> | 0.023 |
| Random intercept effects | Variance | Standard deviation |  |  |
| Lay group | 0.29 | 0.54 |  |  |
| Drop number | 0.001 | 0.03 |  |  |
| Drop order | 0.02 | 0.14 |  |  |
| Drop group | 0.12 | 0.34 |  |  |
| Vial : drop number | 0.03 | 0.18 |  |  |
| Strain replicate : age | 0.03 | 0.17 |  |  |
| Strain replicate : age : sex | 0.05 | 0.21 |  |  |
| Residual | 0.38 | 0.62 |  |  |

### Follow-up experiment testing parental age effects

Throughout the previous experiment, we observed a visible effect of “lay group” on negative geotaxis performance such that vials of flies collected from lay groups 2 and 3 had markedly lower performance than those of lay group 1 (Figure S1). To our knowledge, the only difference between lay groups was parental age, as the focal flies from each lay group came from successive laying events by the same parents. Finding a strong effect of parental age at such relatively young ages (only 1 to 3 days since eclosion) is surprising, so we carried out a third experiment on a more highly replicated subset of strains to see if this parental age effect was repeatable.

Because this experiment was designed primarily to follow-up on our previous results, we followed a nearly identical protocol as that described for the main experiment on the mitonuclear strains, but on a much reduced set of strains to allow for greater replication of each strain at each parental age. In this experiment, we focused on one nuclear background (MAD) that was observed to have high enough reproductive output over time to allow for investigation of parents up to 10 days old. We focused on six mitochondrial strains (ALS, BAR, DAH, HAW, ISR, and JAP) in their replicate populations, for a total of 12 strain replicates. From each strain replicate, we collected 10 pairs of virgin flies, split them into two vials of 5 pairs each, and transferred them to new media every 24 hours for 10 days. After each daily transfer of these parental flies into new vials, we took photos of the eggs laid on the surface of the old vial using the camera of an iPhone 6 to allow us to count total eggs laid over time, prior to trimming eggs from any vials with more than ∼80 eggs, as previously. As offspring began to eclose, we collected 45 virgin males and 45 virgin females from both vials per day of laying per strain replicate; we housed these flies in groups of 15 same-sex individuals, so we collected a total of three 15-individual vials for testing, within each sex, strain replicate, and parental age group. This experiment took place in June 2019, so all strain replicates had also undergone at least 12 additional generations of introgressive backcrossing (for a total of 27 generations for the MAD and ZIM nuclear background lines and 24 for the PUE lines).

Our design therefore resulted in ten lay groups of vials for testing, each differing in parental age by one day. We tested focal flies only at 5 days post-eclosion, transferring them to fresh media once (at 3 days) before testing on day 5. We ran negative geotaxis trials in the Drop Tower on lay groups that had parental ages 1-6, 8, and 10 days. To simplify the scoring of performance from video, we focused on measuring the proportion of flies that climbed off the bottom vial during each drop event rather than calculating the exact average height climbed of all flies in that vial, as this measurement was sufficient to capture variation in our main experiment data set (Figure S1). We paused each video exactly four seconds after vial impact and counted the flies climbing in each vial, then used the number of flies known to be present in each vial to calculate proportion climbing. All flies were tested across five repeated drops, as above.

Further, we used the photos of eggs laid to test whether changing fecundity with parental age relates to differences in offspring performance. Working within ImageJ, we manually counted all visible eggs photographed in every vial. We had egg photographs from two vials of five pairs of parental flies for each strain replicate. We first used this data to test whether fecundity varied across parental age and/or strain. Then, to test how parental fecundity might relate to the negative geotaxis performance of the offspring, we paired each negative geotaxis measurement with the average numbers of eggs laid in the vials from which the focal flies had been sourced.

We performed two different statistical analyses of the data from this experiment. Firstly, we used lme4 to run a linear mixed model testing for effects of parental age and mitochondrial strain on numbers of eggs laid (which followed an approximately normal distribution, validated with a Shapiro-Wilk normality test); this model comprised a response variable of numbers of eggs laid per female, fixed effects of mitochondrial strain (6 levels), parental age (10 levels), and their interaction, and a random slope term allowing the effect parental age to vary across strain replicate. As this model resulted in a singularity warning, we then ran a model with the random slope term simplified to a random intercept of the interaction of parental age and strain replicate. We again used the anova function within the lmerTest package to estimate the significance of fixed effects through F tests using Satterthwaite approximation of degrees of freedom.

Secondly, we ran a generalized linear mixed model with a binomial error distribution to examine the effects of parental age, mitochondrial strain, and sex on the proportion of flies that climbed during the assay. The response variable of this model is a vector of the number of flies climbing during a trial and the number of flies that did not climb. The full model comprised: fixed effects of mitochondrial strain (6 levels), sex (2 levels), parental age (8 levels), their possible two-way interactions, and the average number of eggs in the vials from which the focal flies were collected; random intercept effects of specific group of vials dropped together (drop group, see above) and drop order; a random slope term allowing the effect of drop number to vary across individual vials to account for repeated measures, and a second random slope term to allow the effects of sex and parental age to vary across mitochondrial strain replicates. This maximal model produced a singularity warning, so we reduced the complexity of the random effects statements as described above, and also dropped the most non-significant fixed effects (mitochondrial strain, two-way interaction of mitochondrial strain and sex), until we reached final model that converged on estimates and resulted in no singularity. This final model included: fixed effects of sex, parental age, average number of eggs, the two-way interactions between parental age and sex or mitochondrial strain; and, random intercept effects of drop group, and a composite term that combines mitochondrial strain replicate, sex, and lay group to account for the structure in our data while reducing model term complexity. We evaluated the overall significance of fixed effects of this final model using the Anova function from the *car* package (v. 3.1-3), using type III Wald chi-square tests.

## RESULTS

### Preliminary study on *w^1118^* fly lines

Our results from our preliminary tests on our 13-haplotype set of *w^1118^* strains reveal that male flies had significantly higher negative geotaxis performance than did females, and that performance decreased with increasing fly age (both p < 0.001; Figure 2; Table 2); this age-releated decrease in performance varied significantly across mitochondrial strains (p = 0.03; Figure 2; Table 2).

**Figure 2.**
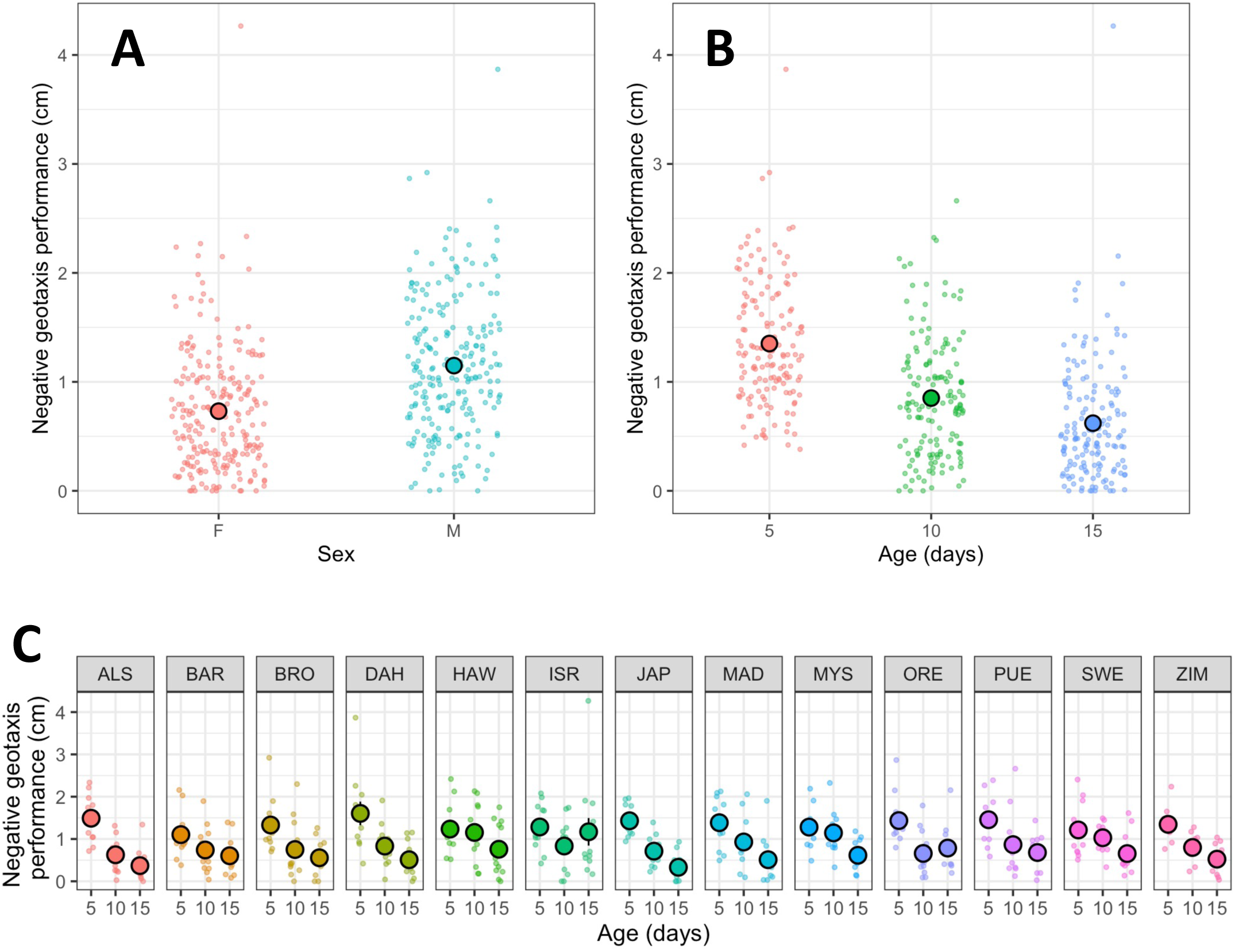
In flies of the *w1118* nuclear background, males (“M”) outperformed females (“F”; A), though performance across all flies decreased with age (B). This age-related decline varied across mitochondrial strains (C). Background points represent the average height climbed (cm) by flies in one vial in one day of testing. Larger points (black outline) represent mean ± SE.

### Main experiment on mitonuclear strains

When we examined negative geotaxis performance across a set of strains comprising 13 mitochondrial haplotypes expressed across three nuclear backgrounds, we found significant effects of the age of the focal flies as well as two-way interactions between nuclear background and sex, nuclear background and age, and sex and age (Table 3, Figure 3, Figure S2). Interestingly, while increasing age tended to decrease negative geotaxis performance generally across all flies, different combinations of sex and nuclear background showed different patterns; for example, while male flies decreased performance with age across all three nuclear backgrounds, only females of the PUE nuclear background demonstrated a marked decrease in performance with age (Table 3, Figure S3). Further, in contrast to the results in our preliminary study in w^1118^ flies, male flies tended to have lower negative geotaxis performance than did females, though the scope of this difference varied with age and nuclear background (Table 3, Figure 3, Figure S2, Figure S3).

**Figure 3.**
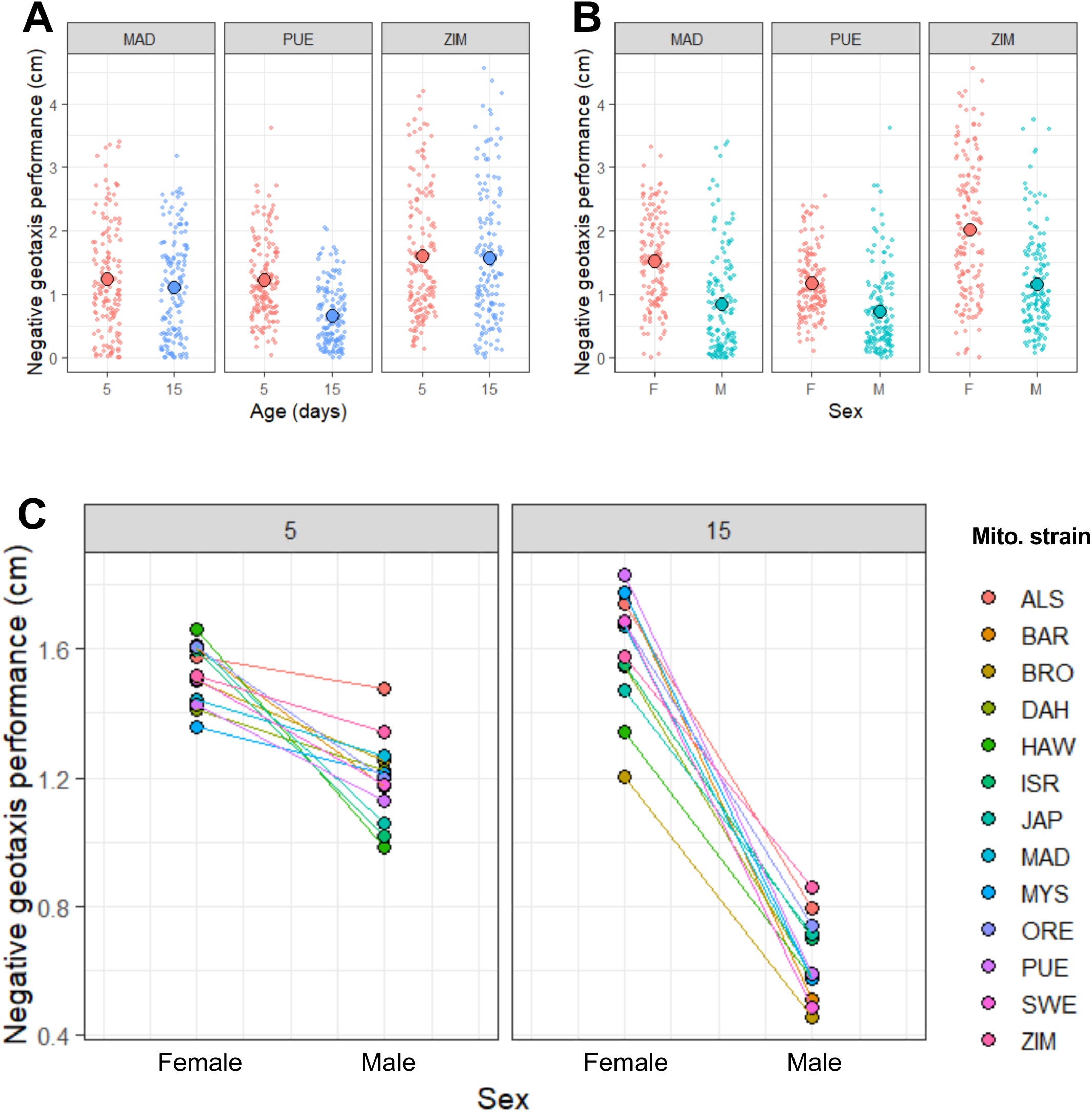
Across the mitonuclear strains of the main experiment, nuclear background significantly affected how negative geotaxis performance varied with age (A) and sex (B), while mitochondrial strain altered how the differences in performance between males and females changed with age (5 or 15 days post-ecolosion; C).

While we found no significant effect of mitochondrial strain alone, the three-way interaction between mitochondrial strain, sex, and age was statistically significant (p = 0.02; Table 3). When we examine this interaction, it appears that younger males (5 days post-eclosion) varied considerably in performance across mitochondrial strains, and the difference in performance between males and females at this age also varied with mitochondrial strain (Figure 3). However, at the older of the two age classes (15 days post-eclosion), females of every strain outperformed males of that strain by a considerable margin; in some strains, females appear to show little to no age-related decline in performance at all (Figure 3, Figure S3). One possible interpretation of this three-way interaction is that while males and females differed more in negative geotaxis performance at day 15 than at day 5, this difference between the sexes varied more across mitochondrial strains at day 5 than at day 15, with males showing especially wide range in performance (Figure 3).

While this experiment was not intended to test for effects of parental age on performance, we did perform the assays across three different “lay groups” (i.e. parental ages; see above) in order to increase our sample size while minimizing variation across other axes. We were surprised to observe that flies of lay groups 2 and 3 (of parental ages two and three days) tended to perform strikingly poorly compared to flies of lay group 1 (of parental age one day; Figure S1), despite no known sources of variation between the groups except parental age and date of sampling.

### Follow-up experiment testing parental age effects

When we tested a subset of our strains (six mitochondrial haplotypes in the MAD nuclear background) across ten different parental ages, we found that flies did, on average, lay fewer eggs with increasing age (Table 4), though visualization of these results suggests an early peak in fecundity at two days of age (Figure S4). Mitochondrial haplotype did not significantly affect fecundity, nor its change with age (Table 4).

**Table 4.** The results of a linear mixed effects model testing the effects of parental age, mitochondrial strain, and their interaction on numbers of eggs laid over 24 hours. We collected these data from a subset of our total strains: six replicated mitchondrial haplotypes in one nuclear background (MAD). We counted eggs from parents laying across 10 days. Subscript values on the F statistic refer to the numerator and denominator degrees of freedom, respectively, approximated with Satterthwaite’s method.

| Fixed effects | Sum of squares | Mean square | F<sub>num, den</sub> | P |
| --- | --- | --- | --- | --- |
| Mitochondrial strain | 142.79 | 28.56 | 0.69<sub>5,108</sub> | 0.63 |
| Parental age | 1329.06 | 1329.06 | 31.94<sub>1,108</sub> | <0.001 |
| Mitochondrial strain : parental age | 84.08 | 16.82 | 0.40<sub>5,108</sub> | 0.85 |
| Random intercept effects | Variance | Standard deviation |  |  |
| Strain replicate : parental age | 7.15 | 2.68 |  |  |
| Residual | 41.60 | 6.45 |  |  |

We found that the negative geotaxis performance of offspring also declined with increasing parental age, and females again outperformed males, as found in the previous experiment (Table 5, Figure 4). This parental-age-related decline in performance does not appear as striking as that found in the previous experiment (Figure S1). However, mitochondrial strain significantly affected the change in performance with parental age (p < 0.001; Figure 4); for example, flies harboring the BAR mitochondrial haplotype appear to show little decrease in performance with increasing parental age, while flies with HAW or JAP haplotypes appear to have a greater tendency toward poor climbing performance at higher parental ages (Figure 4). In addition, negative geotaxis performance in males tended to incur a steeper decline with increasing parental age compared to performance in females, though this effect did not reach statistical significance (p = 0.07; Figure S5).

**Figure 4.**
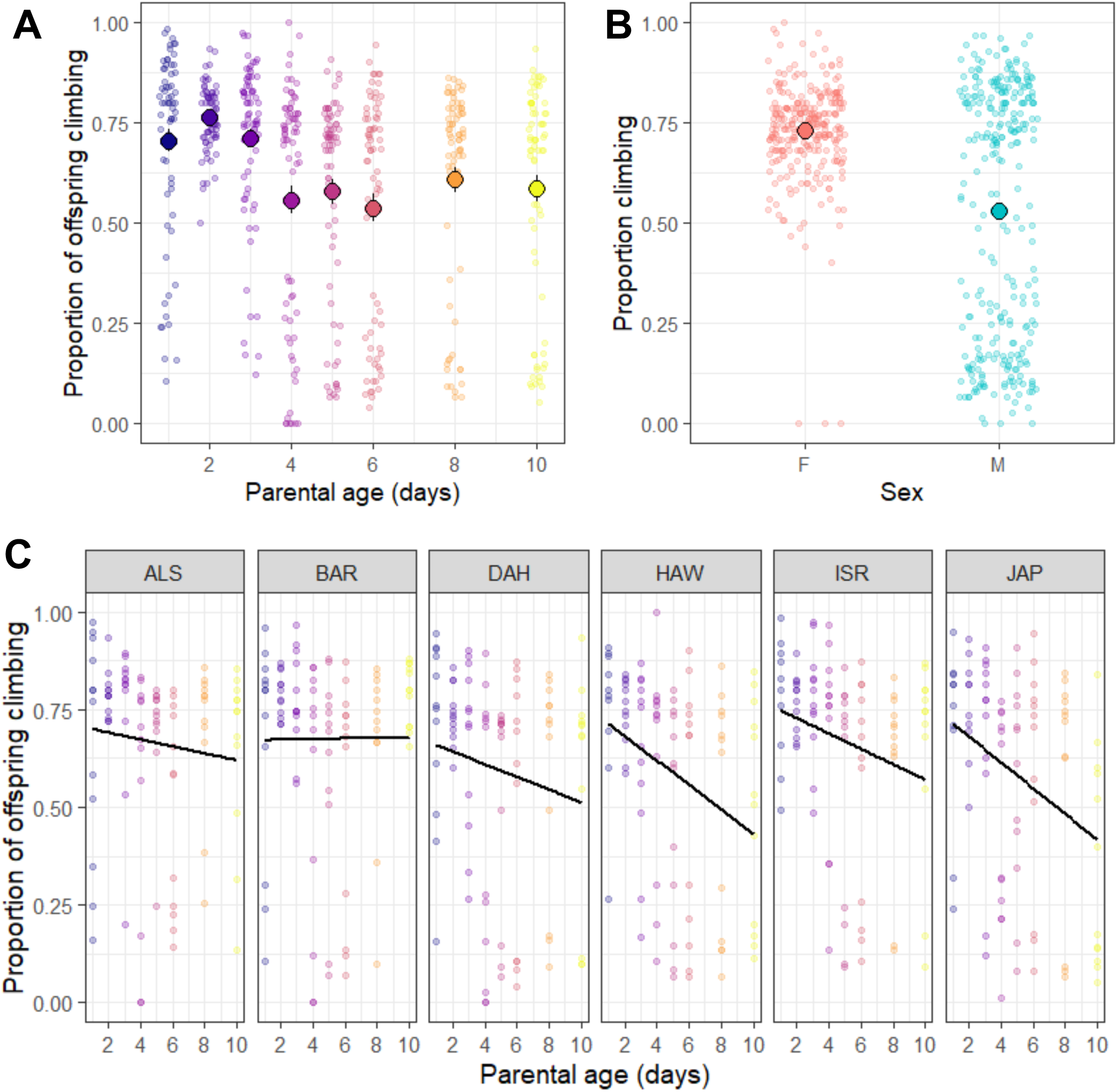
In a follow-up experiment on a subset of strains from the MAD nuclear background, parental age (A) sex (B) affected negative geotaxis performance, here measured as proportion climbing. The effect of parental age on performance varied across flies from different mitochondrial strains (C).

**Table 5.** The results of a generalized mixed effects model (binomial error distribution) testing for the effects of sex, parental age, numbers of eggs laid, and several two-way interactions on negative geotaxis performance. Here, we estimated performance as the proportion of flies that climbed during a trial. We collected these data from a subset of our total strains: six replicated mitchondrial strains in one nuclear background (MAD). All flies tested in this experiment were aged 5 days post-eclosion.

| Fixed effects | Chi-squared | Df | P |
| --- | --- | --- | --- |
| Intercept | 46.40 | 1 | <0.001 |
| Sex | 7.86 | 1 | 0.005 |
| Parental age | 10.59 | 1 | 0.001 |
| Average eggs laid in vial | 16.40 | 1 | <0.001 |
| Sex : parental age | 3.45 | 1 | 0.07 |
| Mitochondrial strain : parental age | 31.8 | 5 | <0.001 |
| Random intercept effects | Variance | Standard deviation |  |
| Drop group | 0.54 | 0.73 |  |
| Strain replicate, sex, and parental age composite term | 0.53 | 0.73 |  |

## DISCUSSION

In this study, we created a new set of 39 replicated mitonuclear strains of *D. melanogaster* fruit flies and tested their negative geotaxis performance, an estimate of locomotory performance. We leveraged these strains in addition to well-studied strains harboring the *w^1118^* nuclear background to examine the relative contributions of mitochondrial, nuclear, and mitonuclear genotypes to negative geotaxis phenotype, along with the effects of age, sex, and parental age. We found that nuclear—but not mitonuclear—genetic variation significantly affected how negative geotaxis performance changed with age and sex in our intraspecific strains of flies. Moreover, while mitochondrial strain alone did not significantly alter performance, mitochondrial genetic variation interacted with either age (in strains carrying the *w^1118^* nuclear background) or both sex and age (in the newly created strains) to affect negative geotaxis phenotype. Our follow-up experiment focusing on a subset of strains in one nuclear background also found that focal fly negative geotaxis performance decreased with increasing parental age, and that this effect differed across mitochondrial strains and might be more pronounced in males.

In contrast to previous studies across mitonuclear strains of *Drosophila* (e.g. Sujkowski *et al*., 2019; Carnegie *et al*., 2021), we found no direct effect of mitochondrial genetic variation on negative geotaxis performance, and we found no evidence for a significant interaction between mitochondrial strain and nuclear background. Instead, across the large set of replicated mitonuclear strains tested in our main experiment, nuclear background shaped the effects of sex or age on negative geotaxis, and mitochondrial strain in turn affected differences between the sexes in age-related decline in performance. Interestingly, the patterns in this interaction between sex, age, and mitochondrial strain suggest a possible Mother’s Curse effect that is detectable only at the younger of the two age classes: at 5 days post-eclosion, males appear more variable in negative geotaxis performance across mitochondrial strains than do females, which is consistent with the “weak form” of Mother’s Curse (Dowling & Adrian, 2019). A study of metabolic rate across these same mitochondrial strains in the *w^1118^*nuclear background found a similar pattern of greater phenotypic variation among strains in males than in females (Nagarajan-Radha *et al*., 2020). Our study did not detect any other markers suggestive of Mother’s Curse, such as evidence of sexually antagonistic mutations or sex-specific mitonuclear interactions (Dowling & Adrian, 2019). However, the results of all three of our experiments indicate that mitochondrial genetic variation can have function effects on how negative geotaxis performance changes with age, sex, and/or parental age.

Recently, two other studies have explored negative geotaxis performance in intra- and inter-specific mitonuclear strains of *Drosophila* fruit flies, focusing on the effects of exercise training via the “Power Tower,” an automated system that repeatedly induces negative geotaxis responses to stimulate flies to climb in a treadmill-like fashion (Piazza *et al*., 2009; Tinkerhess *et al*., 2012a; Sujkowski *et al*., 2019; Spierer *et al*., 2021). These studies found that negative geotaxis performance and exercise response was sensitive to both mitochondrial and mitonuclear genetic variation, though the directions of effects were not always in line with expectations (e.g. some interspecific mixes of mitochondrial haplotype and nuclear background performed better than expectated; Sujkowski *et al*., 2019; Spierer *et al*., 2021). However, in comparison to our current study of both sexes across a large intraspecific set of strains, these studies were performed only on male flies and focused primarily on exercise training response across comparatively divergent mitonuclear strains.

Interestingly, both of these previous studies found that flies with the *w^1118^* nuclear background tended to be relatively insensitive to exercise training and have poor baseline performance (Sujkowski *et al*., 2019; Spierer *et al*., 2021). While negative geotaxis performance of *w^1118^*flies in our preliminary experiment (Figure 2) was within the range of performances of flies with MAD, PUE, or ZIM nuclear backgrounds from our main experiment (Figure 3), we found that males outperformed females only in strains with the *w^1118^*nuclear background, with the opposite pattern in all other backgrounds. Collectively, it appears that *w^1118^* strains tend to respond differently to tests of negative geotaxis than do strains with other nuclear backgrounds; while the genetic and physiological basis for this difference between *w^1118^* and other strains remains undiscovered, *w^1118^* flies have been found to have increased age-related declines in cardiac performance relative to other strains (Kaushik *et al*., 2015), which could plausibly affect their climbing performance. Moreover, the *Drosophila* homologue of the vertebrate gene *PGC1α*, *spargel*, has been found to be key to exercise performance (including negative geotaxis), in part through effects on mitochondria (Tinkerhess *et al*., 2012b; Damschroder *et al*., 2020); exploring strain- and also sex-specific patterns in this and related genes (e.g. Kim *et al*., 2020) is an interesting avenue for further research.

Our findings of a significant effect of parental age also pose a cautionary tale for future studies, given that we detected large variation in performance across sets of flies that were treated near-identically under controlled laboratory conditions, but that differed by 24-48 hours in the age of their parents at time of lay. Based on previous research— including our own preliminary study in the *w^1118^* nuclear background—we did not expect that the relatively small parental age difference (between 1 and 3 days post-eclosion) to have a large effect on offspring performance in our main experiment; parental age effects in *D. melanogaster* are well known, but generally have been studied at much larger parental age differences, such as 10 days or even weeks (e.g. Priest *et al*., 2002; Nystrand & Dowling, 2014; Lee *et al*., 2019). While the serendipitous finding of a strong effect of parental age on offspring performance led to its own experiment here, it emphasizes the importance of considering even small differences in parental age in similar studies (e.g. by varying the ages of the grand- or great-grandparental generation to standardize parental ages); left uncontrolled, these factors could confound experimental designs seeking to partition genetic variance for phenotypic traits.

Overall, we encourage additional exploration of large sets of mitonuclear strains to further validate whether significant effects of mitochondrial strain on phenotype are repeatable across nuclear contexts. While previous studies in the *w^1118^* nuclear background have provided key evidence of male-biased variation across mitochondrial strains and significant effects of mitochondrial haplotype on performance (e.g. Camus *et al*., 2012; Camus & Dowling, 2018; Nagarajan-Radha *et al*., 2020; Spierer *et al*., 2021), other studies of intra- and intersexual sets of strains have found little evidence for Mother’s Curse effects, and have revealed complex mitonuclear interactions (e.g. Mossman *et al*., 2016b; a; Sujkowski *et al*., 2019). Only with further study of diverse sets of strains and traits can we begin to draw broader conclusions about the prevalence and significance of functional mitochondrial variants and Mother’s Curse effects even within this well-studied *Drosophila* system. With the widespread study of negative geotaxis across different lines of research in fruit flies and recent advances in high-throughput methods to quantify performance, such as the Drop Tower described here and the recently developed FreeClimber platform for automating video analyses (Spierer *et al*., 2021), negative geotaxis performance may offer a particularly fruitful avenue for future research into the functional and potentially sex-biased effects of mtDNA variation on locomotion, age-related motor decline, and related physiological processes.

## ACKNOWLEDGEMENTS

This work was supported by the Australian Research Council (DP170100165 and DP200100892 to D.K.D. and DE190100831 to R.E.K.).

