## Supplemental Figures S1-S5 for "Mitochondrial genotype interacts with age and sex, but not nuclear background, to shape locomotory performance across mitonuclear strains of fruit flies"

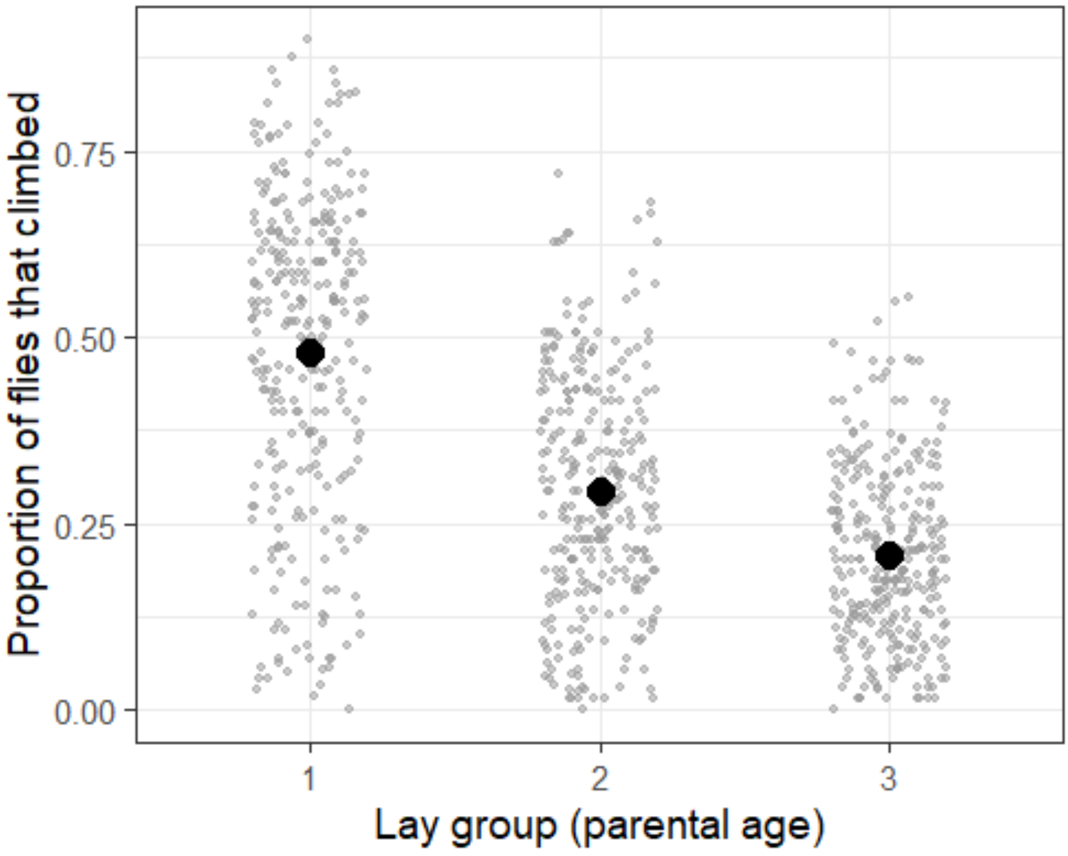


**Figure S1.** Lay group, or parental age (1-2, 2-3, or 3-4 days post-eclosion), appeared to have a strong effect on the proportion of focal flies climbing during a trial in the main experiment.

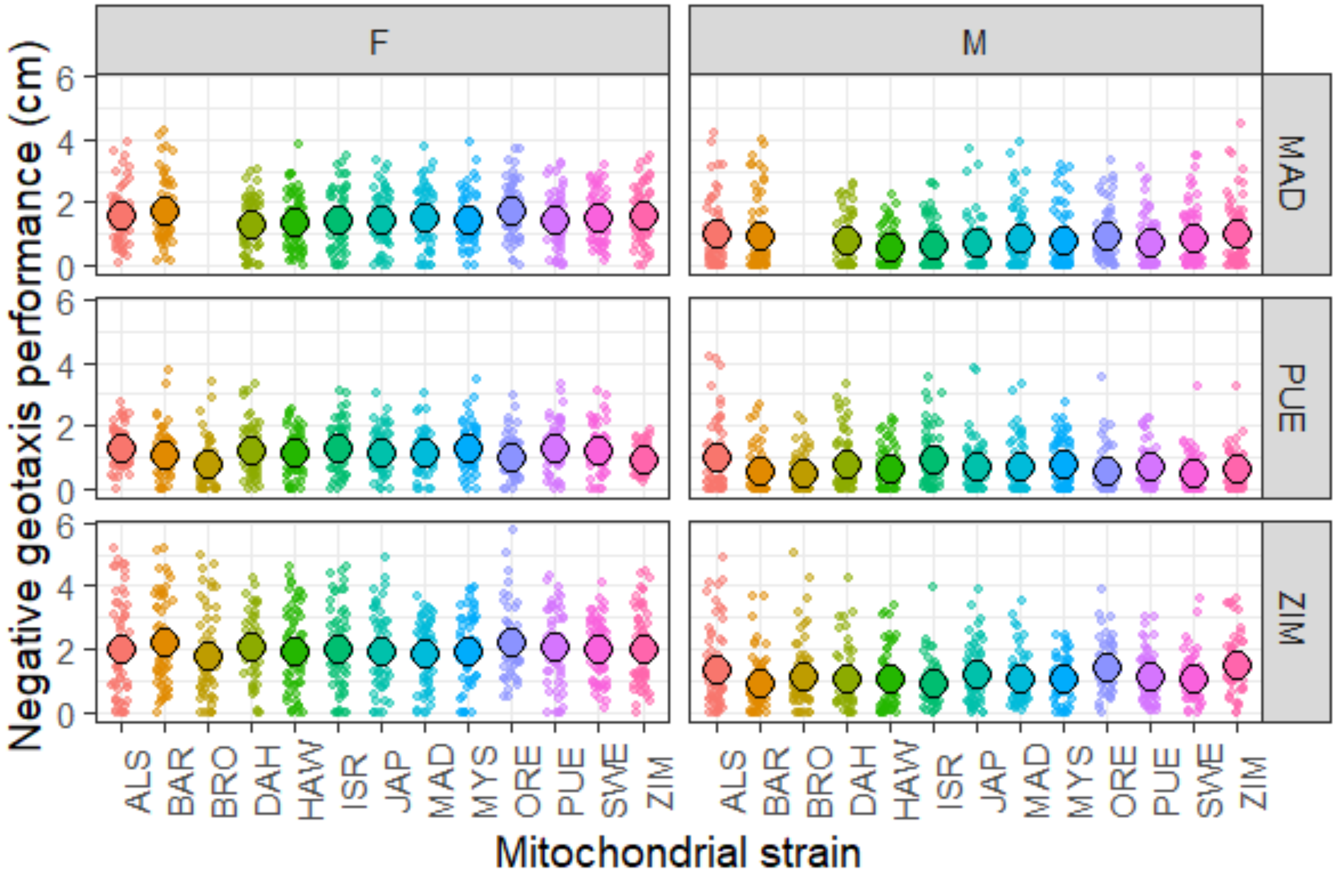


**Figure S2:** Negative geotaxis performance across the 13 mitochondrial strains, three nuclear backgrounds (MAD, PUE, and ZIM), and two sexes (F and M) in the main experiment. Note that these data comprise measures of both strain replicates within each mitochondrial haplotype, and measures at ages 5 and 15 days post-eclosion.

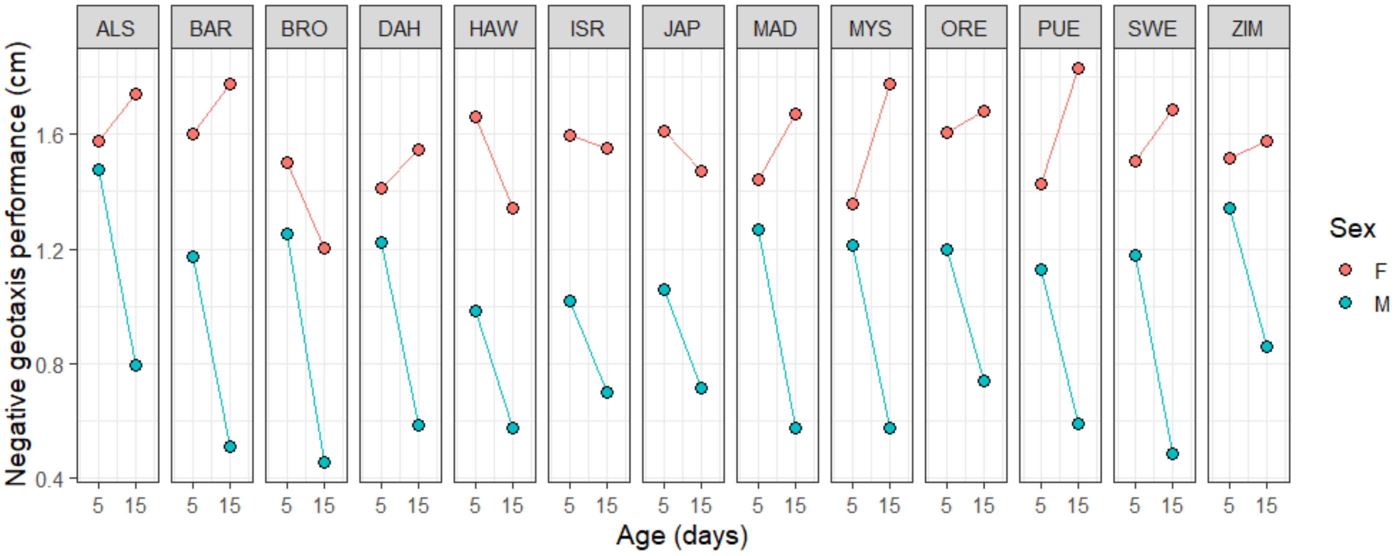


**Figure S3:** The change in negative geotaxis performance with increasing age varied by sex across the 13 mitochondrial strains in the main experiment.

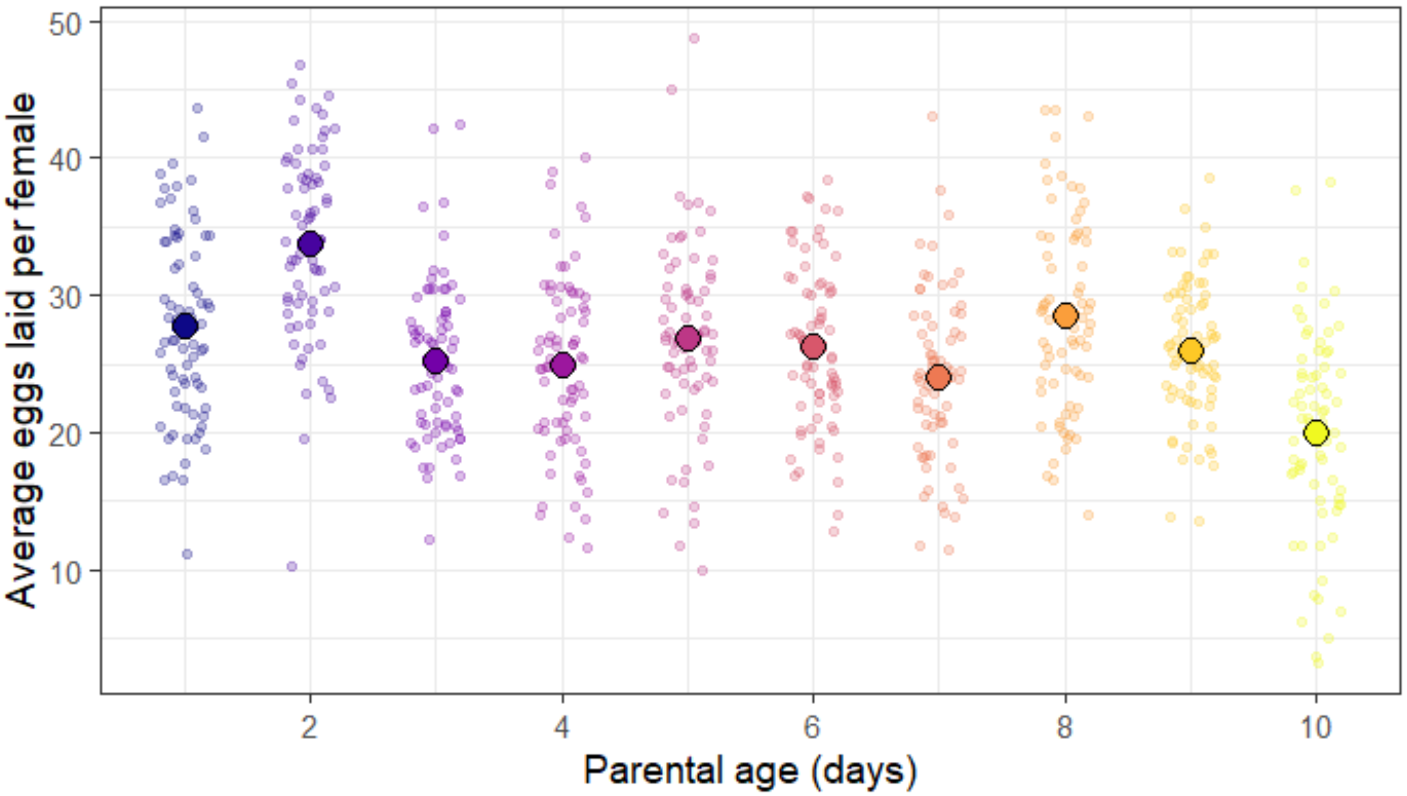


**Figure S4.** Numbers of eggs laid per female decreased with increasing parental age in the third experiment.

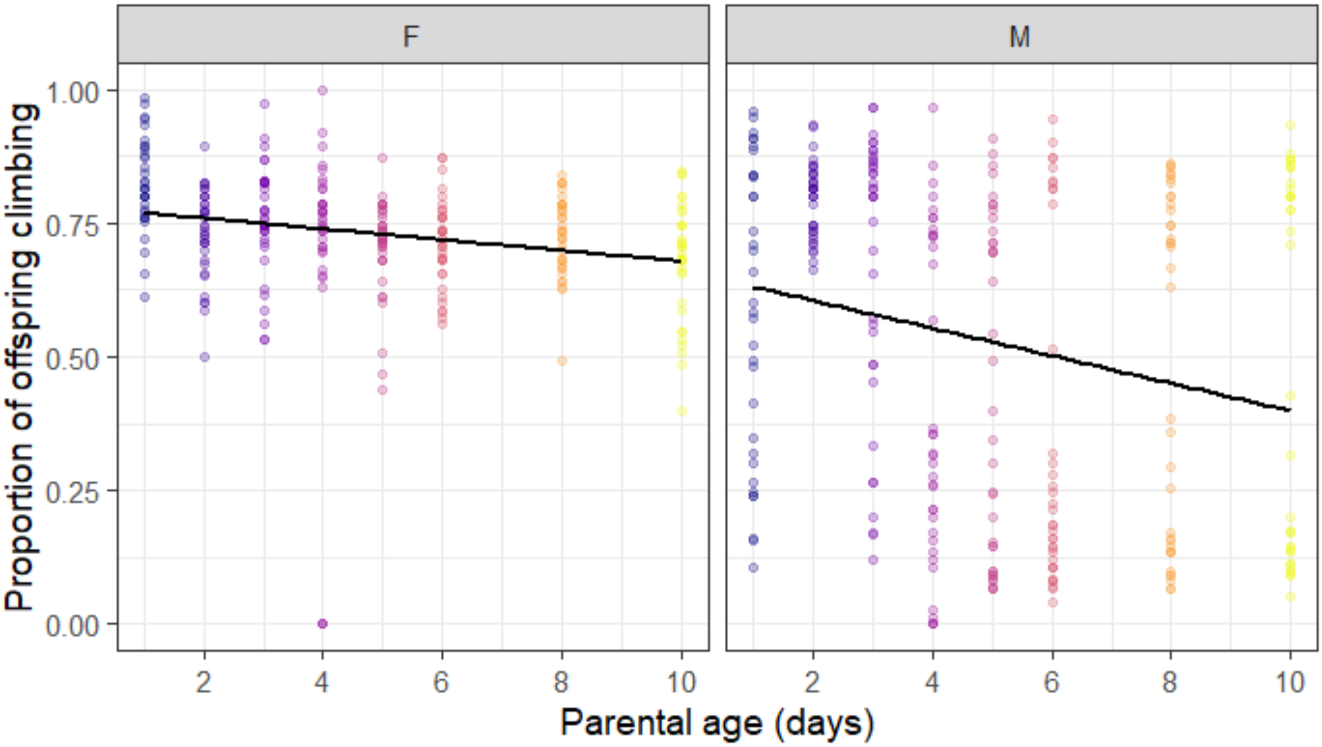


**Figure S5.** Sex weakly affected how negative geotaxis performance changed with parental age in the third experiment, though this effect did not reach statistical significance (p = 0.07).
